# Discovery of high affinity and specificity stapled peptide Bcl-xL inhibitors using bacterial surface display

**DOI:** 10.1101/2023.12.05.570108

**Authors:** Marshall Case, Jordan Vinh, Anna Kopp, Matthew Smith, Greg Thurber

## Abstract

Intracellular protein-protein interactions are involved in many different diseases, making them prime targets for therapeutic intervention. Several diseases are characterized by their overexpression of Bcl-x_L_, an anti-apoptotic B cell lymphoma 2 (Bcl-2) protein expressed on mitochondrial membranes. Bcl-x_L_ overexpression inhibits apoptosis, and selective inhibition of Bcl-x_L_ has the potential to increase cancer cell death while leaving healthy cells comparatively less affected. However, high homology between Bcl-x_L_ and other Bcl-2 proteins has made it difficult to selectively inhibit this interaction by small molecule drugs. We engineered stapled peptides, a chemical modification that can improve cell penetration, protease stability, and conformational stability, towards the selective inhibition of Bcl-xL. To accomplish this task, we built a focused combinatorial mutagenesis library of peptide variants on the bacterial cell surface, used copper catalyzed click chemistry to form stapled peptides, and sorted the library for high binding to Bcl-x_L_ and minimal binding towards other Bcl-2 proteins. We characterized the sequence and staple placement trends that governed specificity and identified molecules with ∼10 nM affinity to Bcl-x_L_ and greater than 100-fold selectivity versus other Bcl-2 family members on and off the cell surface. We confirmed the mechanism of action of these peptides is consistent with apoptosis biology through mitochondrial outer membrane depolarization assays (MOMP). Overall, high affinity (10 nM K_d_) and high specificity (100-fold selectivity) peptides were developed to target the Bcl-x_L_ protein. These results demonstrate that stapled alpha helical peptides are promising candidates for the specific treatment of cancers driven by Bcl-2 dysregulation.

## Introduction

Bcl-2 family members are involved with the regulation of apoptosis, or programmed cell death, via their transient or constitutive interactions with mitochondria. Mitochondria are primarily involved in oxidative phosphorylation and energy production, but these organelles also play an essential role in governing apoptosis.^1,2^ When limits on homeostasis of the cell are exceeded, the cell initiates a complex series of biochemical pathways to initiate programmed cell death. As one of the hallmarks of cancer, cells can evade the protective mechanism of apoptosis and allow the continued survival of the cancer cell.

Central to cancer’s ability to escape apoptosis is blocking the ‘intrinsic’ apoptotic pathway, a carefully regulated signaling pathway of cytosolic and mitochondrial proteins.^3^ On the outer membrane of mitochondria, a signaling network regulates whether mitochondrial membranes remain intact or start large pore formation, releasing cytochrome c and initiating the formation of the apoptosome and eventual cell death.^4,5^ At the center of this pathway, B cell lymphoma 2 (Bcl-2) proteins are interacting with a balance of pro- and anti-apoptotic factors under healthy conditions. However, many cancers overexpress these proteins, which dysregulates mitochondrial function and inhibits apoptosome formation.^6^ The inhibition of Bcl-2 proteins via small molecules or peptides is therefore a direct way to re-establish apoptosis controls for dysregulated cells, providing a powerful approach for the treatment of cancer.^7^

While the Bcl-2 proteins are highly homologous and share many functions, structural differences lead to profound variation in how each of the 5 Bcl-2 proteins (Mcl-1, Bfl-1, Bcl-x_L,_, Bcl-w, and Bcl-2) contribute to apoptosis among other biochemical phenomena.^8,9^ Pathological inquiries have shown that cancers typically overexpress a subset of these 5 proteins. Therefore, selective inhibition of anti-apoptotic proteins has been a longstanding goal of the drug discovery field towards the minimization of off-target toxicity.^6^ However, the design of selective Bcl-2 inhibitors is impactful beyond therapeutic molecules: highly specific inhibitors can be used as ‘tool’ molecules to probe Bcl-2 dependency for novel cancers or used in other biochemical assays. Therefore, discovery of highly specific novel agents may accelerate pathological analyses.^10^ In particular, Bcl-x_L_ remains one of the most important targets among the Bcl-2 family owing to its link with drug resistance, angiogenesis, and cancer cell stemness.^11,12^ Bcl-x_L_ upregulation in breast, glioblastoma, melanoma, among many others is correlated with cancer cell invasion and metastasis.^13–15^ Specific inhibition of Bcl-x_L_ is suggested to address the acquired resistance to poly-specific small molecule drugs like venetoclax.^6,16–20^ However, currently no small molecule drug targeting Bcl-x_L_ has progressed through clinical trials.^6^

Despite the promise of cancer treatment via Bcl-x_L_ inhibition, the discovery of high affinity and selective drugs is challenging for several reasons. First, these proteins rely on alpha helical binding “BH” motifs for recognition of apoptotic proteins, which are shallow and hydrophilic and therefore challenging to target with small molecule drugs.^4,21^ The lack of a traditional hydrophobic binding pocket has led to generation of peptidomimetic drugs which possess many shared characteristics with the proteins the cell naturally uses.^22^ However, the small size of these peptidomimetic drugs often limits their specificity between Bcl-2 members, leading to excessive off-target toxicity.^6^ ABT-737 and its orally bioavailable analog ABT-263 (Navitoclax) were among the most specific small molecule drugs, having high affinity for only Bcl-x_L,_ Bcl-w, and Bcl-2.^21^ However, these drugs resulted in dose limiting thrombocytopenia due to Bcl-x_L_ related toxicity in circulating platelets. Small molecule drugs specifically targeting Bcl-x_L_ have thus far failed in the clinic due to high *in vivo* toxicity. More recently, A-1155463 and A-1331852, small molecules engineered through structure-based drug discovery, have been engineered for high specificity for Bcl-xL over Bcl-2, though they suffer from nanomolar binding to Bcl-w (or were not characterized for Bcl-w binding).^23,24^ To overcome the challenge of targeting large hydrophilic protein-protein interaction domains while mitigating off-target toxicity, scientists have proposed peptides as therapeutics, resulting in high affinity and in some cases high specificity.^12,25–35^. The larger size of peptides enables high affinity and specificity interactions with otherwise difficult to target disease related proteins.^36^ Keating and co-authors sorted a library of linear peptides using yeast surface display, trained machine learning models, and optimized sequences with integer linear programming to achieve high specificities between Bfl-1, Mcl-1, and Bcl-x_L_.^34,37^ Dutta et al. sorted a library of linear peptide variants based on a non-specific but high affinity wild type sequence (BIM), which yielded peptides with ∼1000x specificities between Bcl-xL, Bcl-w, and Bcl-2.^12^ These developments represent important milestones in the development of drug-like Bcl-2 inhibitory peptides. However, additional modifications are needed for drug leads, since linear peptides suffer in the clinic from short *in vivo* half-lives and an inability to penetrate cell membranes.^38–40^ Stapled peptide therapeutics, formed by crosslinking two amino acids, can help address some of the limitations of linear peptides by improving stability, affinity, and membrane permeability.^36,41–50^ However, the development of stapled peptides is typically constrained by low-throughput solid phase peptide synthesis, a bottleneck that limits evaluation of stapled peptides on the order of dozens.^41^

In this work, we use stabilized peptide engineering by *E. coli* display (SPEED) to rapidly evaluate high affinity and specificity Bcl-x_L_ stapled peptides **(Figure 1)**.^51,52^ SPEED enables high-throughput screening of fully stapled peptides in a quantitative format by displaying genetically encoded stapled peptides on the surface of bacteria using non-natural amino acids, enabling evaluation of up to 10^9^ peptides. To generate specific Bcl-x_L_ inhibitors, we designed a library of BIM mutants, a naturally occurring peptide that has high affinity for all Bcl-2 proteins but no specificity towards Bcl-x_L_.^8^ SPEED was used to simultaneously vary both peptide sequence and staple location to determine how staple location governs specificity in the context of a BIM-based library.^51^ We sorted the library using a combination of magnetically activated- and fluorescently activated-cell sorting towards highly specific mutations that would target Bcl-x_L_ with both high affinity and specificity. By analyzing the final peptides in our library using next generation sequencing, we identified sequence trends and staple locations that govern specificity. We then translated select lead compounds off the bacterial surface and show that the discovered peptides retain their properties from surface display in both biolayer interferometry and competitive inhibition experiments.

**Figure 1:**
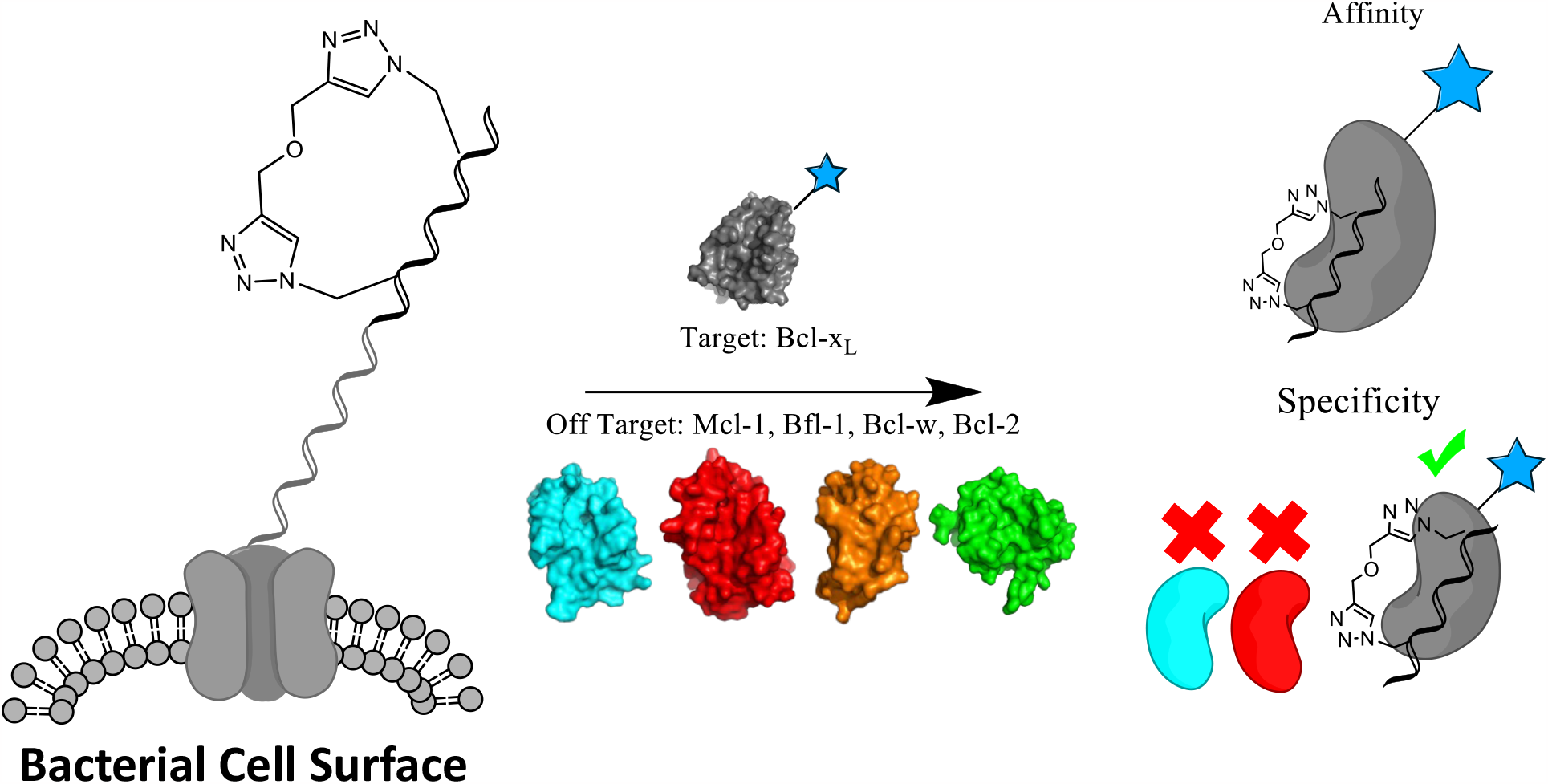
The use of Stabilized Peptide Engineering by *E. coli* Display (SPEED) enables the engineering of high-affinity and -specificity peptides for Bcl-xL.

## Methods

### Purification of Bcl-2 proteins

Bcl-2 proteins were purified as described previously with modifications.^51^ Protein was expressed in E. coli Rosetta2 cells at 37 °C, 250 rpm in terrific broth media supplemented 100 mg/L ampicillin. The purified protein was digested with TEV protease overnight at 4 °C to remove the purification tag in the presence of the following buffer 50mM Tris, pH 7.5, 150mM NaCl, 0.1% β-ME.

### Library Design

To design the library of BIM variants, BH3 sequences and their affinities were collected from literature.^25–27,33,34,53^ Sequences were discretized into 5 bins according to their affinities: <1nM, 1-10nM, 10-100nM, 1000nM. A position specific scoring matrix (PSSM) was generated for each of the 5 Bcl-2 proteins (Bcl-x_L_, Bfl-1, Mcl-1, Bcl-2, or Bcl-w) based on the subset of sequences that had reported affinities for that target. To bias the library design towards high affinity clones, a weighting was applied based on the discretized bins where sequences were counted 1-10,000X in logarithmically spaced bins, depending on which bin it appeared in (high affinity sequences were counted more).

Another PSSM for each Bcl-2 protein was generated by extracting the weights from BIM variant SPOT arrays.^29,30^ First, a binary mask was generated based on the SPOT arrays to separate the location of peptide signal from background. Then, the average intensity of each SPOT was extracted using Fiji v2.0. For each Bcl-2 protein, the two PSSM’s were averaged to capture information from both entire peptide sequences and BIM mutations. Then, mutations were selected to maximize the amount of specificity-driving residues (ones that were highly scored in one library and very weakly scored in others). We excluded Bcl-w and Bcl-2 mutations as they were qualitatively similar to Bcl-x_L_ and we wanted to maintain the balance between Mcl-1, Bfl-1, and Bcl-x_L_ residues. Because bacterial surface display libraries can only be generated on available equipment with ∼10^9^ unique sequences, we constrained the design by first locking amino acids in positions that had the highest absolute magnitude, such as Leu^3a^ and Asp^3f^. Next, we fixed amino acids that didn’t contribute to specificity based on their low weights in the PSSM, such as Gly^1e^, Arg^1f^, Tyr^4d,4e^, and Ala^4f^. Still, the library had more mutations than were possible to display on bacteria. We next eliminated all Cys and Met residues from analysis because we wanted to initially minimize potential disulfide bond issues and out-of-position stapling residues respectively. To put a hard constraint on the size of the library, we merged the processed PSSM’s for Mcl-1, Bfl-1, and Bcl-xL into one by applying the following equation:

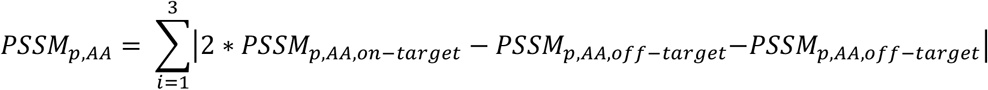

Where p is the position of the peptide sequence, AA is the amino acid, the on-target refers to the PSSM corresponding to the protein target, and the two off-target terms are for the other two Bcl-2 proteins. Finally, degenerate codons were optimized using SwiftLib for a given protein sequence diversity of 10^8^.^54^ The primers and degenerate codon are tabulated in Tables S1 and S2. The design overview is available in Figure S1.

### Library construction

Primers for focused BIM mutants with degenerate codons from the library design step were ordered from IDT whose sequences can be found in Table S3. Libraries were displayed using the pqe80L-eCPX2 plasmid as described previously.^51,52^ We added an N-terminal HA tag and a peptide linker to mitigate steric clashes between display and binding measurements as described previously, resulting in NH_2_-eCPX-HA-linker-peptide-COOH.^25^

### Synthesis and preparation of peptides

Bcl-2 peptides were synthesized, stapled, and purified as described previously.^51,52^ Peptide structures, mass spectra, and chromatograms are in Figure S2 - Figure S4.

### Circular Dichroism

Peptides were dissolved in acetonitrile: water at 1:1 v/v at ∼ 0.1 mg/mL and analyzed on a Jasco J-815 Circular Dichroism (CD) Spectrometer at 100nm/min at 25°C. Spectra were baseline corrected and averaged over 3 runs. Alpha helicity was calculated using the BeStSel web server.^55^ Circular dichroism spectra and alpha helicities are in Figure S4.

### Bacterial Surface Display, Flow Cytometry, and Competitive Inhibition Experiments

The preparation of bacterial cells for flow cytometry and FACS was performed as described previously.^51,52^ Briefly, bacteria expressing the eCPX gene were grown in 1mL M9-methionine-ampicillin overnight. Fresh media was inoculated at a 1:20 ratio and grown for 150 minutes at 37°C. Next, the cells were metabolically depleted for 30 min, then induced with 0.5mM IPTG at 22°C for 4hr with 4mg/mL azidohomoalanine. Peptides on the surface of cells were clicked using 100μM propargyl ether at 4°C for 4hrs. Cells were washed once in PBS before incubation in Bcl-2 protein and expression markers overnight on ice. Cells were washed once in PBS before 15 min incubation with secondary antibody if necessary, then resuspended in PBS before analysis or sorting. Flow cytometry was done using an Attune NXT or BioRad Ze5.

Affinities of peptides on bacterial cell surface were measured using 8 logarithmically spaced concentrations of protein in triplicate. Competitive binding inhibition experiments were performed by incubating a fixed concentration of Bcl-2-biotin protein (100nM for Bcl-xL, Bcl-w, and Bcl-2, and 10nM for Bfl-1 and Mcl-1) with BIM-p5, a known binder for all 5 Bcl-2 proteins, with 8 logarithmically spaced concentrations of peptide in triplicate. After several hours (at least two hours) of incubation at 0°C, 1uL of BIM-p5 displaying bacteria were added for 15 min to capture any unbound Bcl-2 protein. Then the media containing protein-peptide complexes was removed by centrifugation and bacterial cells were prepared for flow cytometry as described above.

### Magnetic Activated Cell Sorting (MACS)

The naïve library underwent three total rounds of magnetic sorting (see Table S4 for sorting details). One round of anti-HA MACS was done using anti-HA magnetic beads (Thermo Fisher). Then, cells were subjected to two rounds of sequential binding-based MACS with 100nM Bcl-x_L_ with the goal of selecting clones below the maximum number of cells that can be analyzed via FACS. The sorting progression is described in Figure S5.

### Fluorescent Activated Cell Sorting (FACS)

Libraries underwent four rounds of sequential fluorescent sorting (see Table S4 for sorting details and Figures S6 and S7 for affinity-based and specificity-based representative FACS plots respectively). The first and second rounds were purely based on affinity and used 100nM and 10nM Bcl-x_L_ respectively. The highest 5% of cells expressing and binding were collected. The third and fourth round were done to improve the specificity of the library. The third round used 100nM Bcl-x_L_ and 25nM of each of the four other Bcl-2 proteins in competition (Mcl-1, Bfl-1, Bcl-w, and Bcl-2). The fourth used 10nM Bcl-x_L_ and 25nM of each of the four other Bcl-2 proteins in competition. In the third and fourth round, the top 5% of cells that were positive for Bcl-x_L_binding and expression but negative for the competitive binding (in a third fluorescent channel) were selected. In a parallel sorting scheme, another third round and fourth round were performed that was a negative sort (25nM each Mcl-1, Bfl-1, Bcl-w, and Bcl-2 for round 3) followed by a positive sort (1nM Bcl-x_L_, round 4). Ultimately, we found that the competitive screens yielded more enrichment than this alternative sorting scheme based on NGS analysis and focused on these sorts for downstream analysis. The sorting progression is described in Table S4 and Figure S5.

### Biolayer Interferometry

Biolayer interferometry was performed as described previously.^51,52^ Representative BLI traces are found in Figure 5.

### Illumina Sequencing and Data Processing

Libraries identified for deep sequencing analysis were prepared as described previously.^51^ Sequencing primers and PCR scheme is shown in Figure S8. Forward and reverse reads DNA were merged using NGmerge and aligned using in-house python scripts. ^56^ We filtered out sequences that didn’t match the framework region of the eCPX2 protein and condensed identical peptide sequences into read counts. To account for the differences in total read counts per library, we converted read counts into frequencies by normalizing to the number of reads. Sequence conservation plots were made using the Logomaker Python package.^57^

### Mitochondrial Membrane Depolarization

Mitochondrial outer membrane polarization assays were performed as described previously.^58,59^ In brief, mammalian cells were added to a mixture of MEB buffer, digitonin, voltage dependent fluorophore (JC-1), oligomycin, beta-mercaptoethanol, with either peptide treatment, FCCP (reversible decoupler of oxidative phosphorylation), alamethicin (a peptide that induces irreversible mitochondrial depolarization), or DMSO (negative control). Fluorescence was measured at 545nm excitation/ 590nm emission using a BioTek Synergy H1 plate reader at 32°C at 5 min intervals for 180 minutes. MDA-MB-231 and Mcf7 cells were used to test Bcl-xL dependence in natural cancer cell lines and the B-ALL leukemia cells were used to test Bcl-2 family dependence.^60^ By normalizing peptide data to the negative control (maximum mitochondrial polarization) and the positive control (complete mitochondrial depolarization), it is possible to measure peptides’ ability to drive cellular apoptosis.

## Results

### Library Design and Cell Sorting

Because the sequence space of BH3-like peptides is much larger than the experimental throughput to measure them (the sequence space of BH3 peptides is ∼10^30^ sequences whereas the throughput of bacteria is ∼10^9^), the primary objective of library design was to identify mutations that are likely to impact specificity. To design such peptides, we noted several design criteria: 1) the library must sample both sequence space and staple location simultaneously to evaluate the impact on affinity and specificity, 2) critical binding residues must be preserved to minimize non-functional variants, and 3) select residues that were predicted to improve affinity and specificity should be mutated to sample the library more efficiently versus random mutagenesis.

First, we chose to simultaneously evaluate staple location and sequence. We hypothesized that epistatic interactions between the staple and the peptide sequence might enable high affinities and/or specificities that would be lost if the staple location was fixed. We chose several staple locations that were previously validated to bind Bcl-2 proteins with varying specificities in the context of BIM.^51^ We chose a peptide length of 23 because peripheral residues generally strengthen binding but do not significantly affect specificity.^12^ Next, we aggregated sequence and affinity data from literature to design a library of variants predicted to improve affinity or specificity towards Bcl-2 members while minimizing the number of non-functional variants. ^25–27,29,30,33,34,53^ We included all naturally occurring and engineered BH3 sequences that had been assayed for binding affinities. Data from SPOT arrays, where the change in binding was measured for BIM single mutants, was combined with sequence data to generate a position specific scoring matrix (PSSM) for each Bcl-2 protein. We weighted mutations based on their affinity to each target and then sampled mutations according to their magnitude of specificity (mutations that were predicted to be highly specific towards one or multiple Bcl-2 proteins) until the desired design space was achieved (∼10^8^ sequences). To maximize the number of sequences from our PSSM and minimize the number of suboptimal residues from degeneracy in the codon table, the final DNA sequences were optimized using SwiftLib.^54^ More details about the computational library design are available in the methods section and Figures S1 and Tables 1-3. After transformation into bacteria, the library had 5.5*10^8^ unique peptides **(Figure 2)**.

**Figure 2:**
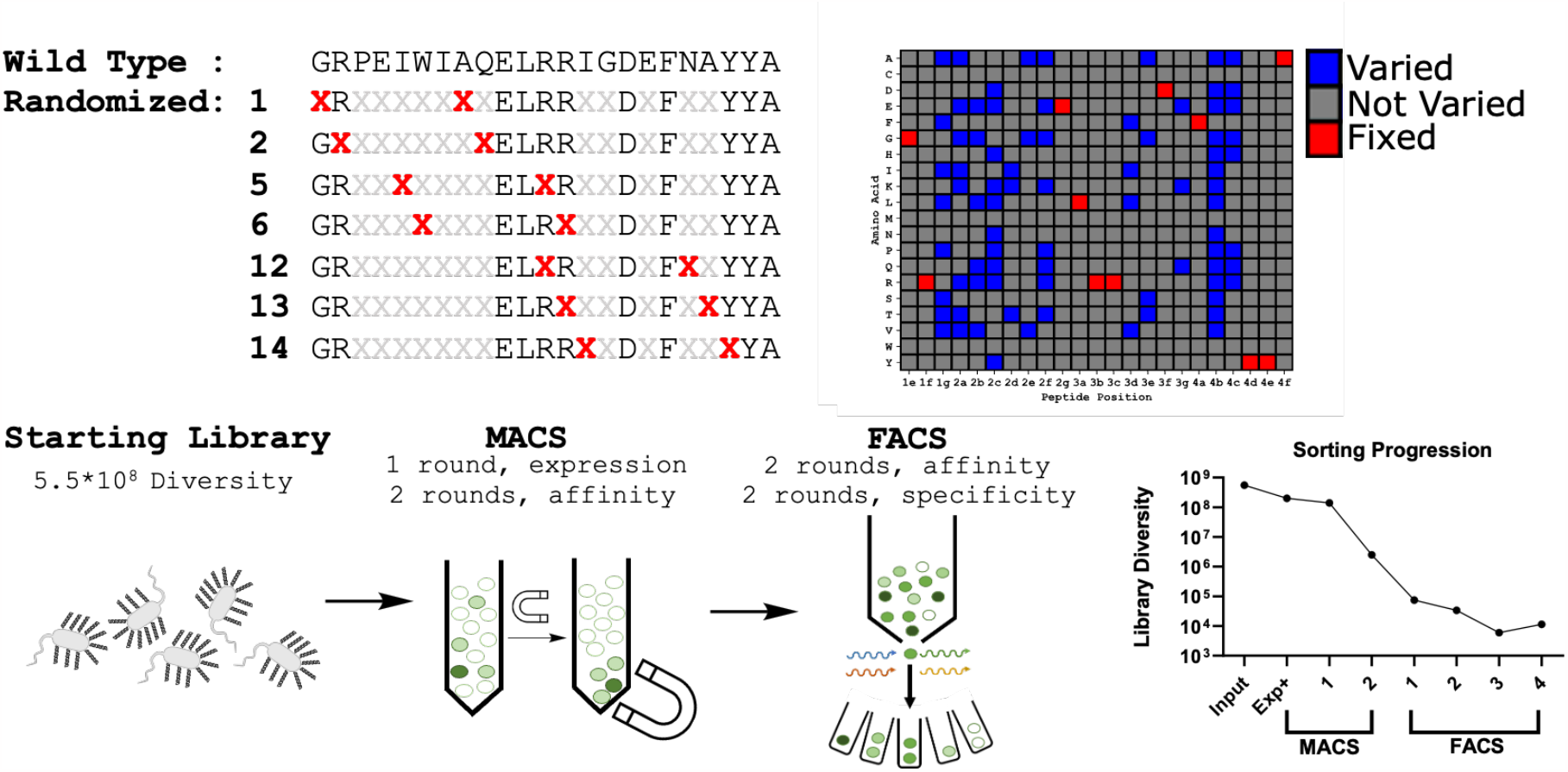
A library of stapled peptides is designed to be enriched with residues and staple positions that govern Bcl-xL affinity and specificity. Staple location and sequence were determined to be two key determinants of affinity and specificity and thus a library where they are simultaneously evaluated was generated. Then, the library is transformed into bacteria, sorted 3 times magnetically and four times fluorescently (in series) before deep sequencing each sort round.

Initial flow cytometry experiments with the naïve library showed only 10-30% of all cells were displaying peptide, indicating that many cells either had ampicillin resistance but did not contain functional plasmid or that peptide-eCPX2 was being inefficiently shuttled to the cell surface. To enrich the libraries towards functional peptides, we performed 3 rounds of magnetic activated cell sorting (MACS) followed by 4 rounds of fluorescent activated sorting (FACS) (see Table S4 for sorting details, Figures S6 and S7 show representative FACS plots, and Figure S9 has logoplots for FACS rounds 1-4). MACS experiments were split into two phases: one round to improve the expression of the library and two to improve binding while simultaneously shrinking the library to a size that could be sorted by FACS, where more specific boundaries can be chosen for desired peptide fitness. FACS experiments were also split into two phases: the first two rounds were designed to improve the affinity of the library towards Bcl-x_L_ by screening with decreasing concentrations of protein (100nM and 10nM respectively), and the last two rounds were done to improve the specificity by performing competition experiments and selecting towards highly specific binders. We hypothesized that this new library would be optimally sorted via FACS by first finding the high affinity Bcl-x_L_ binders and then identifying the subset that were highly specific as previously reported.^27^ After sorting, we performed low throughput flow cytometry experiments, which suggested that the library had been highly enriched towards specific binding, as evidenced by nearly saturable binding at low concentrations (<10nM) of Bcl-x_L_ but minimal binding towards other Bcl-2 proteins even at high concentrations (>100nM). In addition to sorting for highly specific peptides with competition sorting experiments, we also tested whether a round of negative sorting (the lack of binding to off-target proteins) followed by a positive round of sorting (target binding) would yield highly specific clones. Both competitive and non-competitive sorts yielded a similar set of enriched sequences (Figure S10), suggesting the library was well suited to finding specific Bcl-x_L_ inhibitory stapled peptides.

### Next Generation Sequencing

After magnetic and fluorescent sorting, we analyzed the set of enriched peptides along the sorting progression using Illumina NovaSeq next generation sequencing (NGS) **(Figure 3)**. We first investigated how sorting influenced the enrichment and proportions of the library composition (**Figure 3a)**; all rounds of sorting resulted in an enrichment of sequences and depletion of others. We next investigated the relationship between the staple position and the peptide sequence among highly functional Bcl-x_L_ peptides. Despite staple scanning in the context of BIM suggesting that many positions had high affinity or specificity for Bcl-x_L_ (the 2^nd^, 6^th^ and 12-14^th^ staple position BIM mutants had high affinity toward Bcl-x_L_), surprisingly, all stapled positions except the 6^th^ were nearly eliminated after the first round of FACS (**Figure 3b)**.^51^ While most sequences observed had the 6^th^ staple position, peptide sequence is highly dependent on the location of staple (Figure S11). For example, while Lys^2d^ is the dominant mutation for the 6^th^ position, Ile is more prevalent for the 5^th^ and 14^th^ position. This suggests that there exists a complex relationship between the staple and the sequence that supports simultaneous screening of staple location and peptide sequence.

**Figure 3:**
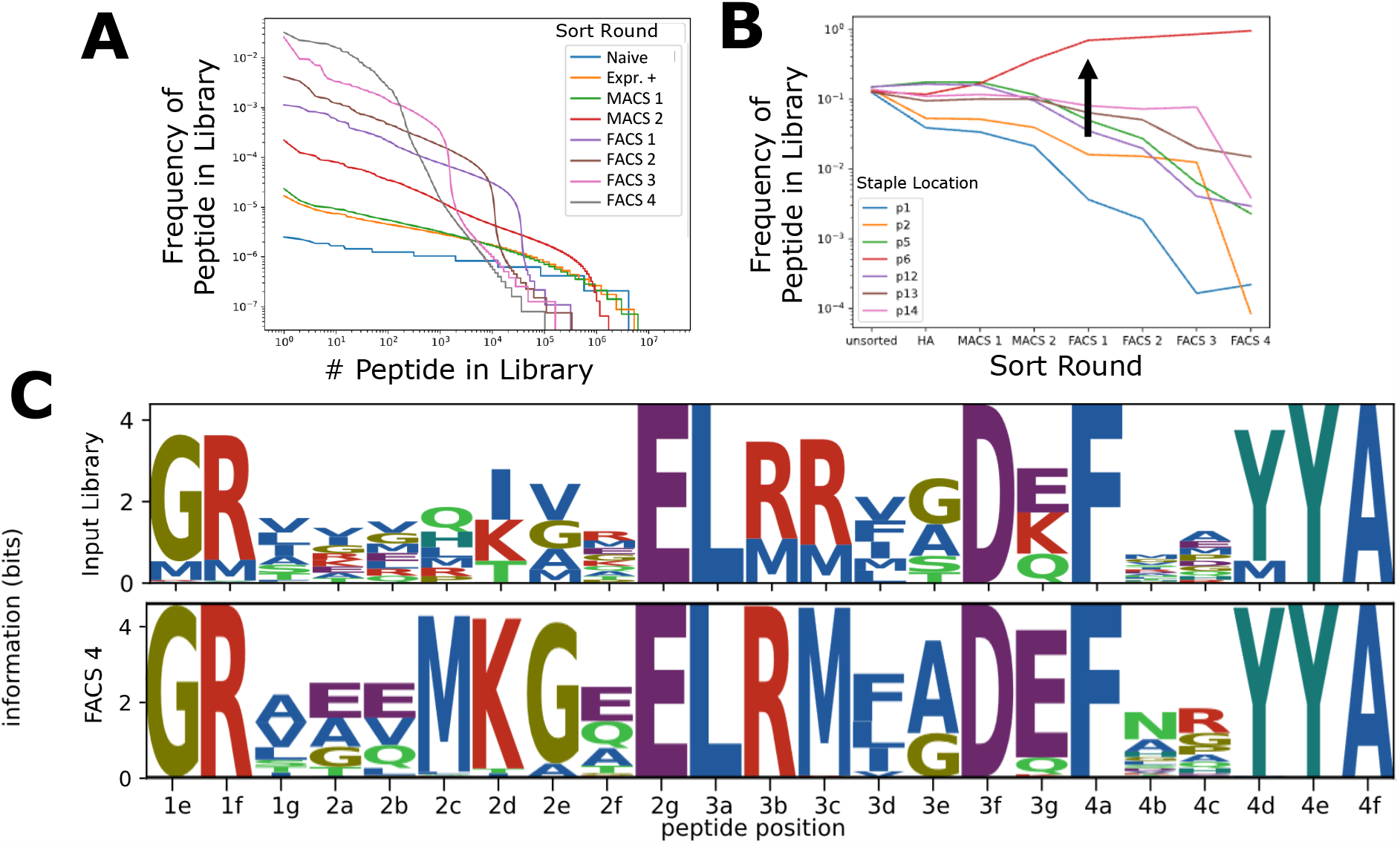
Next generation sequencing of sorted Bcl-xL peptides yields insights into the staple location and sequence patterns that govern specificity. (A) First, the distribution of peptide sequences’ enrichments was calculated across sorting rounds. Whereas there is little bias in the library composition in the naïve library, as indicated by its nearly uniform distribution, each subsequent round of sorting biases the library towards a subset of sequences. By the final rounds of sorting, ∼10^3^ sequences represent nearly 100% of all sequences remaining. (B) By the second round of magnetic sorting, the 6^th^ staple position nearly dominates the library and continues to displace other staple positions, suggesting that this staple position contributes to affinity and specificity. (C) Compared to the naïve library (top), which displays no bias towards certain residues in randomized positions, the Bcl-xL sorted library has clear sequence conservation patterns.

Next, we looked for conserved sequence patterns and whether these matched other Bcl-x_L_ peptides previously described (**Figure 3c)**. Compared to the naïve library, where any position not fixed by design displays nearly uniform distribution between selected mutations, the Bcl-x_L_ library has clearly conserved sequence patterns. The most dominant mutations, Lys^2d^, Gly^2e^, and Asp^3g^, appeared in nearly every peptide that remained in the library. Positions 2a, 2b, and 2f did not display the same level of conservation but generally have enriched negative glutamic residues, whereas the naïve library had mostly small hydrophobic residues. Positions 1g, 4b, and 4c had a smaller magnitude of enrichment, consistent with their position peripheral to the main alpha helical interface.^11^

### Evaluation of peptide hits

The sequencing results and analyses at the library level suggested that the peptides had been selected towards high affinity and specificity Bcl-x_L_ peptides. We next investigated the affinity and specificity for individual sequences. We randomly selected thirteen of the highest frequency clones from the final round of FACS and evaluated their affinity and specificity towards the Bcl-2 proteins **(Figure 4)**. First, we measured binding at two concentrations of target, significantly above (100nM) and below (1nM) the median library binding affinity, to obtain a crude estimate of affinity. First, we measured the binding of each of these peptides on the bacterial cell surface towards Bcl-x_L_ at 1nM in triplicate. This concentration was chosen because it does not saturate BIM-p5 for Bcl-x_L_, which is a double-digit nanomolar binder. Thus, increases in binding when normalized to display level are indicative of an improvement in binding affinity. All sequences evaluated demonstrated significantly improved binding, suggesting that peptide hits had K_d_’s in the low double digit nanomolar range. We similarly evaluated the binding of the other Bcl-2 proteins and observed significantly decreased binding towards Mcl-1 and Bfl-1 (∼10,000 fold weaker binding). However, the magnitude of binding towards Bcl-w and Bcl-2 was not reduced to the same extent (10-100 fold weaker binding), though most clones had statistically significantly reduced binding for all four off-target proteins (Figure S12). We then tested binding at 100nM, which should saturate all but the weakest binders. This analysis suggested that most peptides were highly specific for Bcl-x_L_, though fewer peptides had statistically significant specificity for Bcl-x_L_ relative to Bcl-w and Bcl-2.^12^

**Figure 4:**
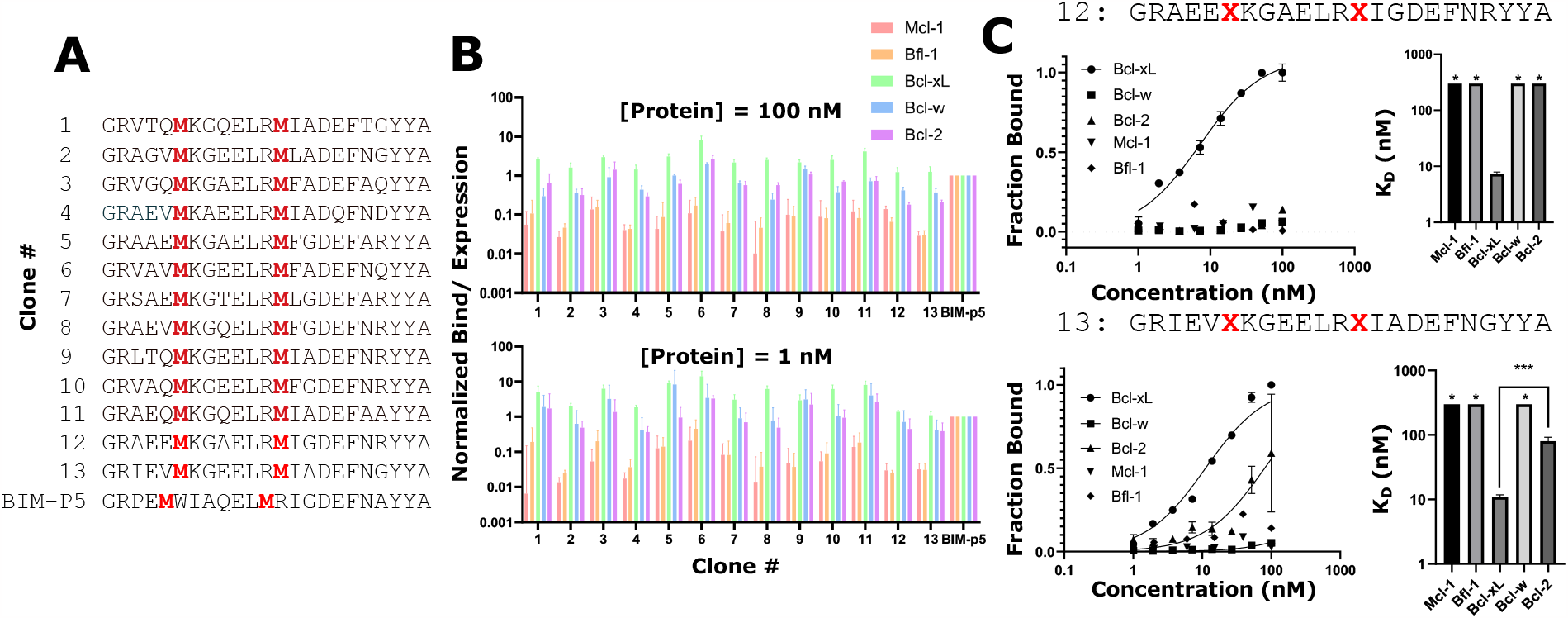
Individual peptides from the library are highly specific towards Bcl-xL. (A) Thirteen peptides were randomly chosen from the final round of cell sorting and their sequences were identified via Sanger sequencing. Each of these peptides were measured for their binding against all 5 members of the Bcl-2 protein family at (A)1nM, which is significantly below the K_d_ of the wild type sequence for the main target, Bcl-xL, or (C) 100nM, which is a high enough concentration to saturate all but the weakest of binders. (D)Two of the thirteen peptides were chosen for more thorough analysis and their binding affinities were measured on the cell surface by flow cytometry.

Based on the promising crude estimates from the library, we selected two clones for further analysis. These variants were selected based on their diminished binding towards all 4 off-target Bcl-2 proteins while maintaining high affinity towards Bcl-x_L_. We titrated two of these clones, denoted **12** and **13** with various concentrations of proteins on the cell surface. These peptides had high affinities towards Bcl-x_L_ (∼10 nM K_d_) and greatly weakened affinity towards the other targets, with the exception of **13** binding to Bcl-2 with ∼80 nM K_d_. The identification of these specific stapled peptides for Bcl-x_L_ further demonstrated the ability of SPEED to find diverse sequences that have desirable activity.

### Solution Phase Measurements

SPEED has been previously validated to predict binding properties of stapled peptides with comparable accuracy to solution phase measurements.^51^ However, to ensure the binding affinities measured via bacterial surface agreed with solution phase peptide binding, we synthesized the two peptides using solid phase peptide synthesis. After stapling the peptides (as detailed in Methods), we characterized the peptides’ secondary structures using circular dichroism (see Figure S4). Both peptides had moderate alpha helicity before stapling (18% and 47%) but had significantly enhanced helicity when stapled (44% and 67%, respectively). Then, we used two techniques to measure the binding affinity and specificity: competitive inhibition and biolayer interferometry **(Figure 5)**. In the competitive inhibition experiments, peptides of various concentrations are equilibrated with select soluble Bcl-2 protein before the addition of BIM-p5 displayed on the surface of *E. coli* bacteria. After a short incubation of equilibrated peptide and protein with bacteria, any unbound protein is rapidly sequestered by bacteria, which are analyzed via flow cytometry. The fraction of protein bound is inversely proportional to the fraction of protein blocked by soluble peptide. The K_i_ is calculated by fitting the fraction bound as the peptide concentration is titrated before converting to a K_d_ based on the affinity of BIM-p5 to soluble protein using the Cheng-Prusoff equation. In the biolayer interferometry experiments, a streptavidin sensor is loaded with biotinylated Bcl-2 protein and incubated with various concentrations of soluble peptide. Real time measurements allow the determination of equilibrium K_d_ and kinetic rates of k_on_ and k_off_ (kinetic parameters and statistics are available in Figure S13). Both solution phase techniques gave similar results to bacterial surface display measurements, in agreement with previous reports.^51,52^ As previous reports confirm that bacterial surface measurements of affinity may overestimate affinity, disagreement between the value of **13** binding Bcl-2 via SPEED versus those from solution phase peptides confirm that solution phase measurements are still recommended. This analysis further demonstrated that both evaluated peptides had specificities towards Bcl-x_L_ over Bcl-w by an order of ∼100, Bcl-2 by ∼1,000, and all others by >10,000.

**Figure 5:**
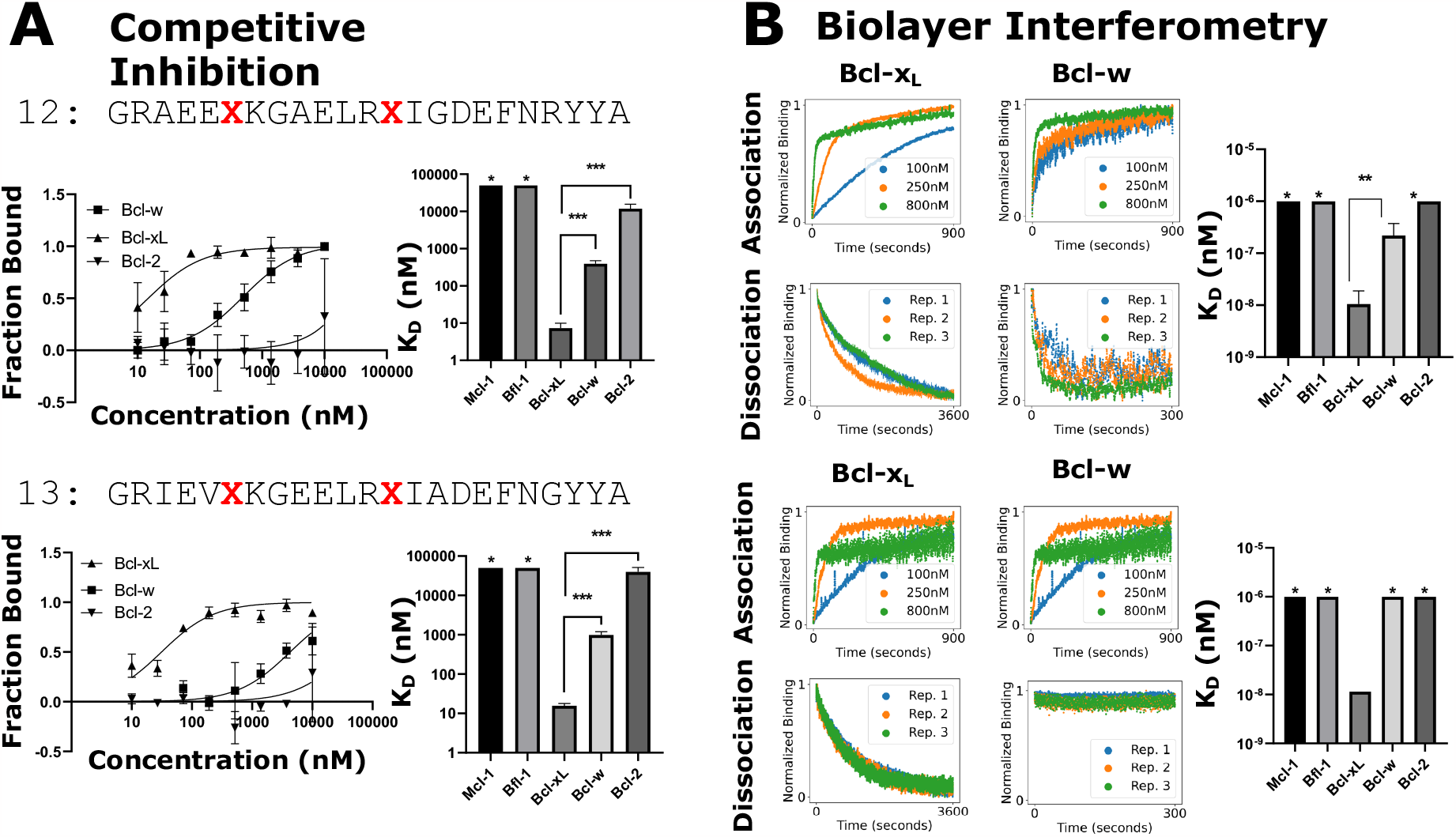
Solution phase characterization of select Bcl-xL stapled peptides. (A) Both hits’ binding affinities are assayed by competitive inhibition. In this experiment, soluble peptide (in excess) and protein are allowed to equilibrate before a high-affinity peptide displayed on bacteria is added, which sequesters any unbound protein. The bacteria are then analyzed via flow cytometry and the fraction bound is inversely proportional to soluble peptide binding. The K_i_’s calculated and converted to a Kd using the Cheng-Prusoff equation. (B)The binding affinity is also measured using biolayer interferometry, where a sensor is loaded with biotinylated protein and the binding of peptide at various concentrations is measured in real time. The on and off rate are fit using GraphPad Prism v10.0. The K_d_ calculated as the ratio of k_off_ to k_on_. *: no binding detected. **: p < 0.01 ***: p < 0.005.

### *In vitro* characterization

We next sought to characterize whether peptides functionally induced apoptosis in human cancer cell lines **(Figure 6)**. On the surface of mitochondria, Bcl-2 proteins sequester Bak and Bax, which hetero-oligomerize to form pores and depolarize mitochondria. This phenomenon can be measured using a voltage sensitive fluorophore (JC-1) through a mitochondrial outer membrane polarization (MOMP) assay.^58,59^ First, we incubated MDA-MB-231 and MCF7 (human derived cancer cells overexpressing Bcl-xL) with various concentrations of **13** with digitonin (to facilitate membrane permeability) and found that the peptide was able to depolarize mitochondria with nanomolar concentrations **(Figure 6)**. Next, we tested a peptide known to not bind Bcl-xL (**F2**, binding affinities are shown in Figure S14) and demonstrated that this peptide was not able to depolarize the cells, indicating specific Bcl-xL binding is required. Finally, we measured the B-ALL cell lines which are engineered to overexpress specific Bcl-2 proteins and showed that **13** did not depolarize cell lines overexpressing other members of the Bcl-2 family **(Figure 6c)**.^60^

**Figure 6:**
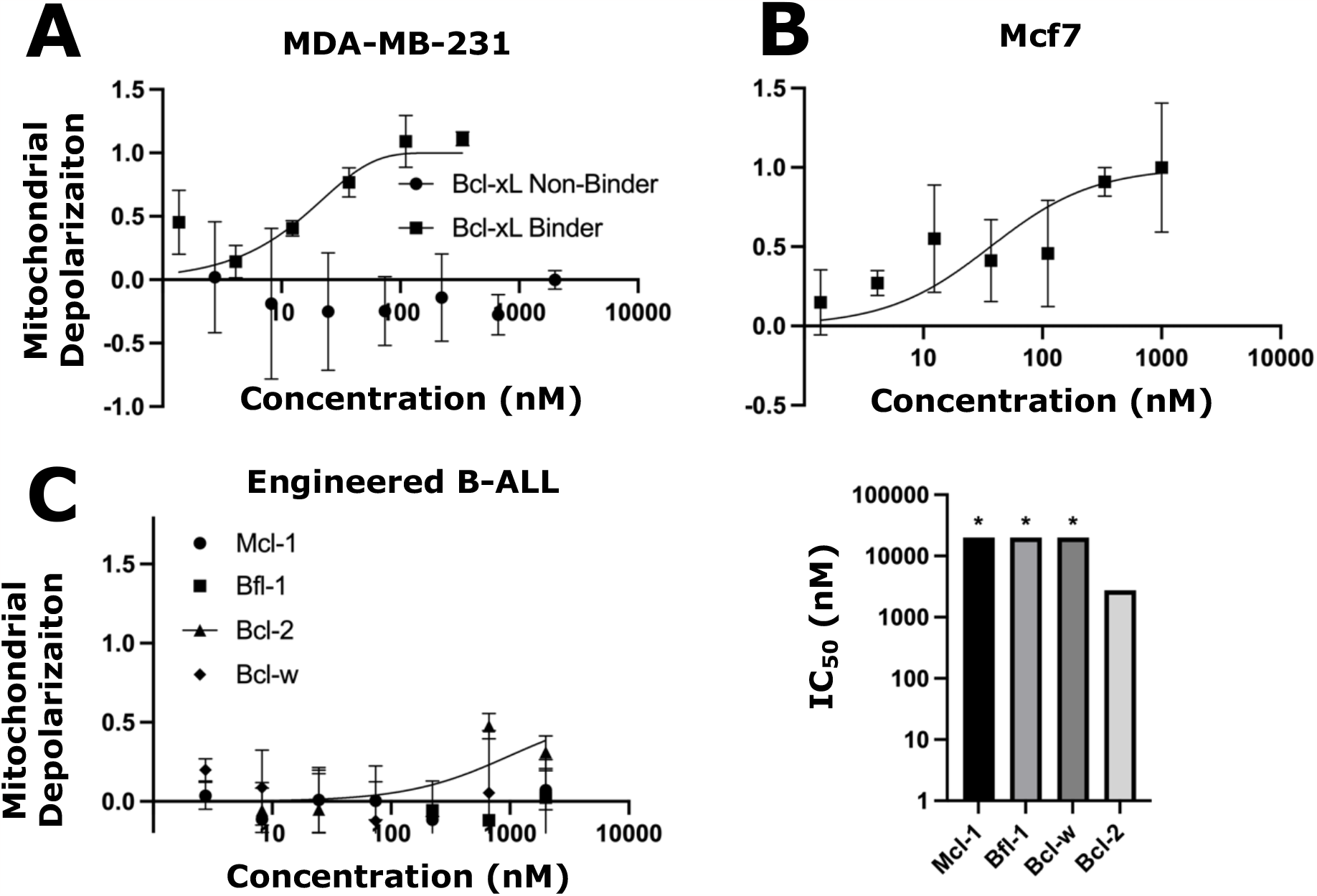
The mechanism of action is confirmed via mitochondrial outer membrane polarization (MOMP). MOMP is measured for compound **12** at various concentrations in two Bcl-xL dependent cell lines: MDA-MB-231 (A) and MCF7 (B). The specificity of the molecule was measured by titrating B-ALL cell lines, engineered to be driven by a specific Bcl-2 protein, with compound **12**. The K_i_ of the peptide was fit using GraphPad Prism v10.0 and is shown (bottom right).

## Discussion

Direct targeting of Bcl-2 proteins is an effective approach to restore apoptosis in cancerous cells. Therefore, high affinity and specificity molecules targeting the B cell lymphoma 2 (Bcl-2) family of proteins have important therapeutic potential. In this work, we used SPEED to screen stapled peptides with varying linker location and sequence to select specific and high affinity compounds towards Bcl-x_L_ **(Figure 1)**. Given the large theoretical sequence space, we designed a focused computational library that we hypothesized contained mutations that would drive specificity between Bcl-xL and the other Bcl-2 proteins **(Figure 2)**. To accomplish this design task, we leveraged two sources of data: sequence-affinity databases and SPOT arrays of BIM mutants. ^25–27,29,30,33,34,53^ First, we pooled the body of BH3 peptides’ sequences and affinities for Bcl-2 proteins. While it has been shown that higher order epistatic effects are present in BH3 peptides, many more mutations act non-epistatically, and thus we assumed independent mutational effects on affinity as a reasonable simplification.^34^ We aggregated these data and used them to predict which mutations were most likely to govern affinity and/or specificity (see Methods). After optimizing degenerate codons that sampled these critical residues and constraining the library size to ∼10^8^, we transformed the library into bacteria, yielding a library with 5.5*10^8^ diversity.

A combination of magnetic (MACS) and fluorescent (FACS) sorting, focusing on affinity or specificity, were used to enrich the library towards specific Bcl-x_L_ peptides **(Figure 3)**. While MACS was used to improve the expression of the library and then enrich the library to a diversity smaller than 10^7^, FACS was the primary tool used to achieve affinity and specificity in the peptide libraries due to its ability to directly select from the binding/expression landscape. We previously established that the staple location is a major driver of specificity.^51^ Because SPEED enables simultaneous evaluation of sequence mutants and staple locations, we were able to sample this complex relationship across the sort progression. Sequence patterns that emerged among highly specific Bcl-x_L_ peptides generally agree with previous reported sequences. For example, we observed that Phe^3d^ was the most highly enriched amino acid and has been suggested to destabilize binding towards Mcl-1.^12,61^ Both Glu^2f^ and Glu^3g^ were previously shown to be specific for Bcl-xL.^26^ Both Phe^3d^ and Leu^3d^ appear more abundantly in the final round than Ile^3d^, which has been suggested to drive specificity towards Mcl-1. However, we also achieved specificity while not requiring many previously established mutations, such as Val^4a^ or Lys^4e^, which drives specificity for Mcl-1 or Bcl-2 respectively but was not randomized in our library (instead, Phe^4a^ and Tyr^4e^).^12,61^ This suggests further improvements could be identified by further combining specificity-driving mutations discovered in this report with those reported elsewhere.

Data from both flow cytometry and next generation sequencing at the library level suggested that peptides were highly specific towards Bcl-x_L._ To confirm that this was true for individual peptides, we sampled sequences randomly from the final round of sorting and evaluated their affinity and specificity compared to a known binder towards all 5 proteins, BIM-p5 **(Figure 4)**.^51^ With significantly more specificity towards Mcl-1 and Bfl-1 than Bcl-2 or Bcl-w, these peptide libraries are consistent with reports that achieving specificity between the three Bcl-proteins is a more challenging task and that further improvements to sorting strategy or new mutations are necessary to achieve higher margins of specificity.^12^ Two peptides with maximum specificity were chosen for a more thorough analysis beyond the two-concentration binding estimates as predictive of affinity. These peptides showed ∼10 nM affinity and > 10-fold specificity for all 4 family members on the bacterial cell surface, demonstrating a successful selection of high affinity and specificity peptides for Bcl-x_L_. The ability of SPEED to evaluate the fitness of a peptide with few flow cytometry samples makes it amenable to analyzing hits quickly before more thorough downstream analysis.

While bacterial cell surface experiments suggested that molecules discovered from sorting were high affinity and strongly specific, it also tends to weakly overestimate affinities, and we therefore sought to translate those molecules into solution phase to confirm their binding properties **(Figure 5)**.^51^ After synthesizing, stapling, and purifying the peptides using standard Fmoc chemistry (see Methods), we measured the binding properties of the peptides using two methods: competitive inhibition and biolayer interferometry. Both methods closely agreed and confirmed SPEED measured affinities; both peptides analyzed were extremely specific, having ∼10nM affinity for Bcl-x_L_ but >200nM K_d_ for Bcl-w and >10,000nM K_d_ for all others. Biolayer interferometry measurements additionally measure kinetic parameters, yielding insight into whether the on- or off-rate dominates the specificity differences. Comparing the binding between Bcl-x_L_ and Bcl-w suggests that off-rates drive specificity (Figure S13); k_on_ for Bcl-w and Bcl-x_L_ are not significantly different (p=0.266) while k_off_ for Bcl-w is more than 40 times faster than that of Bcl-x_L_ (p = 0.00201). These results confirm previous reports that bacterial surface display measured affinities highly correlate with those from solution phase, whether from competitive inhibition experiments or biolayer interferometry. ^51^

We further sought to confirm that the peptides were acting with mechanisms consistent with apoptosis biology. To confirm that the affinities and specificities on or off the bacterial cell surface were not artifacts of soluble versions of the Bcl-2 proteins, and to confirm they act consistently with known apoptosis mechanisms, we measured the permeabilization of mitochondrial outer membranes when titrated with peptides **(Figure 6)**. ^58,59^ Bcl-xL overexpressing cell lines (MDA-MB-231 and MCF7), were depolarized by the Bcl-xL specific peptide (**13**) but not a Bcl-xL non-binder **(Figure 6a**,**b)**. In contrast, cell lines not driven by Bcl-xL (B-ALL engineered for Mcl-1, Bfl-1, Bcl-w, or Bcl-2 overexpression) were not depolarized by the Bcl-xL specific peptide **(Figure 6c)**. These results confirmed that peptides discovered via SPEED act in accordance with apoptosis biology.

While SPEED yielded high affinity and specificity Bcl-x_L_ stapled peptides, there are some limitations and improvements for future work. While flow cytometry experiments confirmed improvements in fitness as sorting progressed, we speculate that our campaign could have been improved by incorporating specificity-based sorting earlier on. High affinity and non-specific peptides were more abundant than highly specific but weakly binding peptides; the elimination of non-specific peptides earlier may have yielded specific hits with fewer rounds of sorting. This would likely be more important in future experiments that target Bcl-w and Bcl-2, where there are fewer defined mutations that improve specificity compared to Bcl-x_L_, Mcl-1, or Bfl-1.

In conclusion, we used SPEED to engineer high affinity and highly specific Bcl-x_L_ stapled peptide antagonists. We demonstrated they are highly specific when presented on the cell surface or synthesized in soluble form, act in accordance with apoptosis biology, and have unique structural motifs that enable their high specificity. The enriched library can be utilized to incorporate design rules towards cell permeability and protease stability to further increase these important drug-like properties.^62,63^ Overall, SPEED can be used as a versatile platform to the generate potent and specific stapled peptides.

