## Supplemental Tables and Figures for "Discovery of high affinity and specificity stapled peptide Bcl-xL inhibitors using bacterial surface display"

Running title:

**Table S1:** Predicted and observed masses of peptides chemically synthesized in this study.

| Compound | Predicted Unstapled Mass | Observed Unstapled Mass | Predicted Stapled Mass | Observed Stapled Mass |
|---|---|---|---|---|
| **12** | 2736.3 | 2736.3 | 2830.4 | 2830.4 |
| **13** | 2681.3 | 2681.3 | 2776.4 | 2776.4 |

**Table S2:** Degenerate codon design for each staple location for the peptide library.

| | 1e | 1f | 1g | 2a | 2b | 2c | 2d | 2e | 2f | 2g | 3a | 3b | 3c | 3d | 3e | 3f | 3g | 4a | 4b | 4c | 4d | 4e | 4f |
|---|---|---|---|---|---|---|---|---|---|---|---|---|---|---|---|---|---|---|---|---|---|---|---|
| **wt** | G<br>ggt | R<br>cgc | P<br>ccg | E<br>gaa | I<br>att | W<br>tgg | I<br>att | A<br>gcg | Q<br>caa | E<br>gaa | L<br>ttg | R<br>cgc | R<br>cgc | I<br>att | G<br>ggt | D<br>gac | E<br>gaa | F<br>ttt | N<br>aac | A<br>gcg | Y<br>tat | Y<br>tat | A<br>gcg |
| **p1** | M<br>atg | R<br>cgc | X<br>dya | X<br>rna | X<br>sda | X<br>can | X<br>aha | M<br>atg | X<br>vva | E<br>gaa | L<br>ttg | R<br>cgc | R<br>cgc | X<br>ntc | X<br>gsa | D<br>gac | X<br>vaa | F<br>ttt | X<br>vnc | X<br>svm | Y<br>tat | Y<br>tat | A<br>gcg |
| **p2** | G<br>ggt | M<br>atg | X<br>dya | X<br>rna | X<br>sda | X<br>can | X<br>aha | X<br>gba | M<br>atg | E<br>gaa | L<br>ttg | R<br>cgc | R<br>cgc | X<br>ntc | X<br>rsc | D<br>gac | X<br>vaa | F<br>ttt | X<br>vnc | X<br>smm | Y<br>tat | Y<br>tat | A<br>gcg |
| **p5** | G<br>ggt | R<br>cgc | X<br>dya | X<br>rna | M<br>atg | X<br>can | X<br>awa | X<br>gba | X<br>vva | E<br>gaa | L<br>ttg | M<br>atg | R<br>cgc | X<br>ntc | X<br>rsc | D<br>gac | X<br>vaa | F<br>ttt | X<br>vnc | X<br>smm | Y<br>tat | Y<br>tat | A<br>gcg |
| **p6** | G<br>ggt | R<br>cgc | X<br>dya | X<br>rna | X<br>sda | M<br>atg | X<br>aha | X<br>gba | X<br>vva | E<br>gaa | L<br>ttg | R<br>cgc | M<br>atg | X<br>ntc | X<br>gsa | D<br>gac | X<br>vaa | F<br>ttt | X<br>vnc | X<br>svm | Y<br>tat | Y<br>tat | A<br>gcg |
| **p12** | G<br>ggt | R<br>cgc | X<br>nyc | X<br>rna | X<br>sda | X<br>cna | X<br>aha | X<br>gba | X<br>vva | E<br>gaa | L<br>ttg | M<br>atg | R<br>cgc | X<br>ntc | X<br>rsc | D<br>gac | X<br>vaa | F<br>ttt | M<br>atg | X<br>smm | Y<br>tat | Y<br>tat | A<br>gcg |
| **p13** | G<br>ggt | R<br>cgc | X<br>dya | X<br>rna | X<br>sda | X<br>cna | X<br>awa | X<br>gba | X<br>sva | E<br>gaa | L<br>ttg | R<br>cgc | M<br>atg | X<br>ntc | X<br>rsc | D<br>gac | X<br>vaa | F<br>ttt | X<br>vnm | M<br>atg | Y<br>tat | Y<br>tat | A<br>gcg |
| **p14** | G<br>ggt | R<br>cgc | X<br>dya | X<br>rna | X<br>sda | X<br>cna | X<br>aha | X<br>gba | X<br>vva | E<br>gaa | L<br>ttg | R<br>cgc | R<br>cgc | M<br>atg | X<br>gsa | D<br>gac | X<br>vaa | F<br>ttt | X<br>vnc | X<br>svm | M<br>atg | Y<br>tat | A<br>gcg |

**Table S3:** Primers used to incorporate peptide library for bacterial surface display.

| Primer Name | DNA Sequence |
|---|---|
| P1 | GCTGGCCAGTCTGGCCAGTATCCGTATGATGTGCCGGATTATGCGGCGGGCGGCAGCGGCGGCAGCGGCGGCCAGAGCATGCGCDYARNASDAVNGAHAATGVNGGAATTGCGCCGCDTSRSCGACVAATTTVNC<br>GMCTATTATGCGGGAGGGCAGTCTGGGCAG |
| P2 | GCTGGCCAGTCTGGCCAGTATCCGTATGATGTGCCGGATTATGCGGCGGGCGGCAGCGGCGGCAGCGGCGGCCAGAGCGGTATGDYARNASDAVNGAHAGBAATGGAATTGCGCCGCDTSRSCGACVAATTTVNC<br>SMMTATTATGCGGGAGGGCAGTCTGGGCAG |
| P5 | GCTGGCCAGTCTGGCCAGTATCCGTATGATGTGCCGGATTATGCGGCGGGCGGCAGCGGCGGCAGCGGCGGCCAGAGCGGTCGCDYARNAATGVNGAHAGBAVVAGAATTGATGCGCDTSGSAGACVAATTTVNC<br>SVMTATTATGCGGGAGGGCAGTCTGGGCAG |
| P6 | GCTGGCCAGTCTGGCCAGTATCCGTATGATGTGCCGGATTATGCGGCGGGCGGCAGCGGCGGCAGCGGCGGCCAGAGCGGTCGCDYARNASDAATGAHAGBAVNGGAATTGCGCATGDTSGSAGACVAATTTVNC<br>GCGTATMNSGCGGGAGGGCAGTCTGGGCAG |
| P12 | GCTGGCCAGTCTGGCCAGTATCCGTATGATGTGCCGGATTATGCGGCGGGCGGCAGCGGCGGCAGCGGCGGCCAGAGCGGTCGCDYARNASDAVNGAHAGBAVNGGAATTGATGCGCDTSGSAGACVAATTTATG<br>GCGTATMNSGCGGGAGGGCAGTCTGGGCAG |
| P13 | GCTGGCCAGTCTGGCCAGTATCCGTATGATGTGCCGGATTATGCGGCGGGCGGCAGCGGCGGCAGCGGCGGCCAGAGCGGTCGCDYARNASDAVNGAHAGBAVNGGAATTGCGCATGDTSGSAGACVAATTTVNC<br>ATGTATTATGCGGGAGGGCAGTCTGGGCAG |
| P14 | GCTGGCCAGTCTGGCCAGTATCCGTATGATGTGCCGGATTATGCGGCGGGCGGCAGCGGCGGCAGCGGCGGCCAGAGCGGTCGCDYARNASDAVNGAHAGBAVNGGAATTGCGCCGCATGRSCGACVAATTTVN<br>MGMCATGTATGCGGGAGGGCAGTCTGGGCAG |
| eCPX reverse | GAGGTCATTACTGGATCTATCAACAGGAGTCCAAGCTCAGC |

**Table S4:** Primary and secondary labeling conditions for all fluorescently activated cell sorting in this study.

| Sort Round | Binding | Expression | Off-Target |
|---|---|---|---|
| **MACS expression** | N/A | 1st: anti-HA magnetic beads | N/A |
| **MACS 1** | 1st: 100nM biotin-target + streptavidin magnetic beads | N/A | N/A |
| **MACS 2** | 1st: 100nM biotin-target + streptavidin magnetic beads | N/A | N/A |
| **FACS 1** | 1st: 100nM biotin-target<br>2nd: 1:100 neutravidin-DL488 | 1st: 1:100 anti-HA mouse<br>2nd: 1:100 chicken anti-mouse-AF647 | N/A |
| **FACS 2** | 1st: 10nM biotin-target<br>2nd: 1:100 neutravidin-DL488 | 1st: 1:100 anti-HA mouse<br>2nd: 1:100 chicken anti-mouse-AF647 | N/A |
| **FACS 3 competitive** | 1st: 100nM AF647-target | 1st: 1:100 anti-HA mouse<br>2nd: 1:100 goat anti-mouse-AF488 | 1st: 25nM biotin-competitors<br>2nd: 1:100 neutravidin-DL405 |
| **FACS 4 competitive** | 1st: 10nM AF647-target | 1st: 1:100 anti-HA mouse<br>2nd: 1:100 goat anti-mouse-AF405 | 1st: 25nM x 4 biotin-competitors<br>2nd: 1:100 neutravidin-DL488 |
| **FACS 3 negative** | N/A | 1st: 1:100 anti-HA mouse<br>2nd: 1:100 goat anti-mouse-AF405 | 1st: 25nM x 4 biotin-competitors<br>2nd: 1:100 neutravidin-DL488 |
| **FACS 4 positive** | 1st: 10nM AF647-target | 1st: 1:100 anti-HA mouse<br>2nd: 1:100 goat anti-mouse-AF405 | N/A |

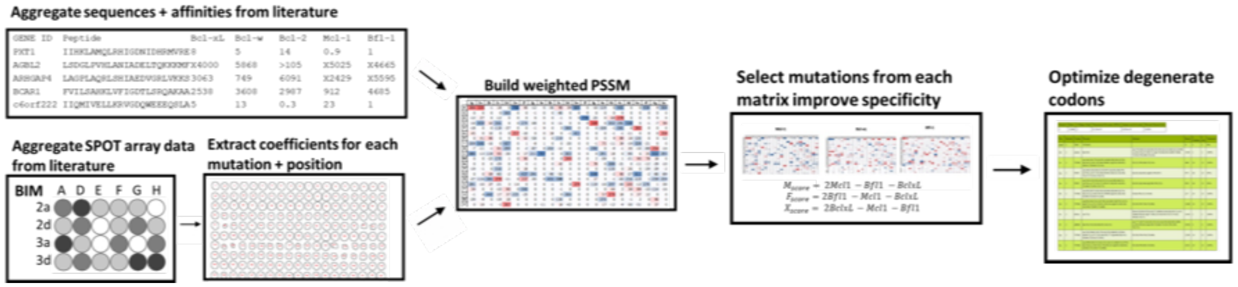


**Figure S1:** Representation of library design scheme. First, BH3-like sequences with measured affinities to one or more Bcl-2 proteins are aggregated and aligned from literature. Sequences are binned according to their affinity ($K_d$ < 1 nM, < 10nM, < 100nM, <1,000nM and > 1,000 nM) and weighted proportionately to the tightness of their binding (10,000X, 1,000X, 100X, 10X and 1x). Separately, the impact of mutations to BIM (measured via SPOT array data) is similarly aggregated. Weights from each spot are extracted using FIJI. Both sequence and spot data are averaged to build a weighted position specific scoring matrix (PSSM). This process is repeated for each Bcl-2 protein (Mcl-1, Bfl-1, etc.) and degenerate codons are designed to maximize the extremeness of a mutation for a particular Bcl-2 protein (i.e. mutations that minimize Mcl-1 but maximize others). See Methods for information on the optimization of degenerate codons.

**12**

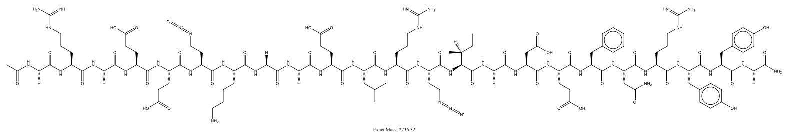


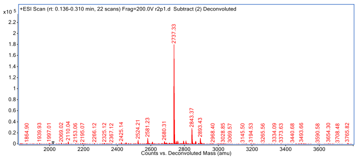


**12-PE**

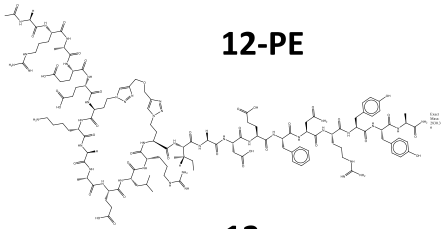

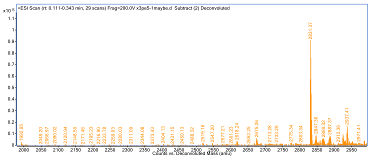


**13**

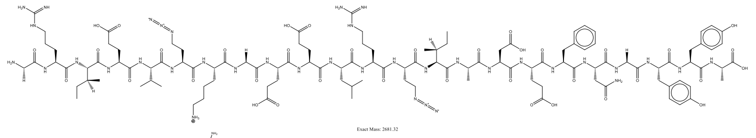


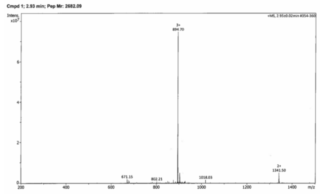


**13-PE**

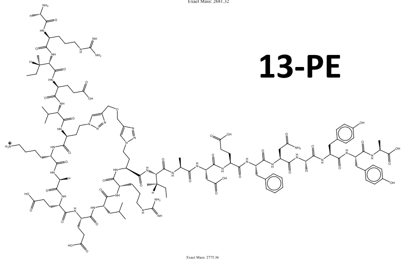

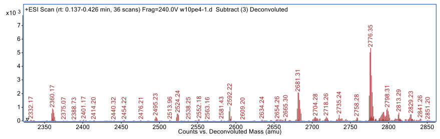


**Figure S2:** Representative structures and experimental mass spectra for all compounds chemically synthesized in this study.

**12**

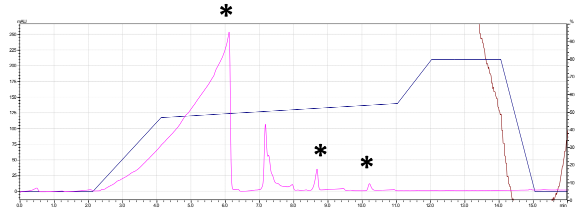

**12-PE**

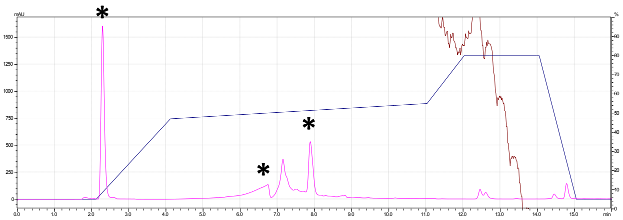

**13**

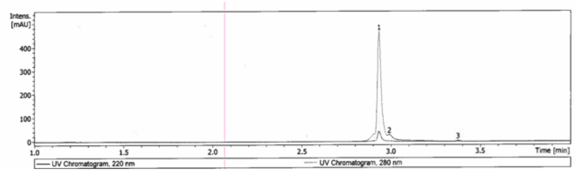


**13-PE**

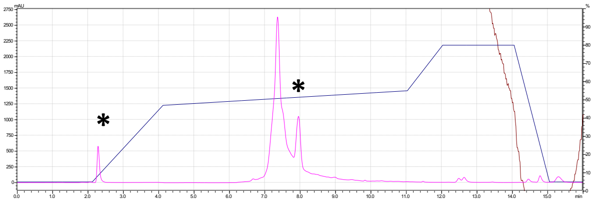

**Figure S3:** High performance liquid chromatography (HPLC) chromatograms for all compounds used in this study. Asterisks (*) denote false peaks due to an issue with the detector unit and do not represent impurities.

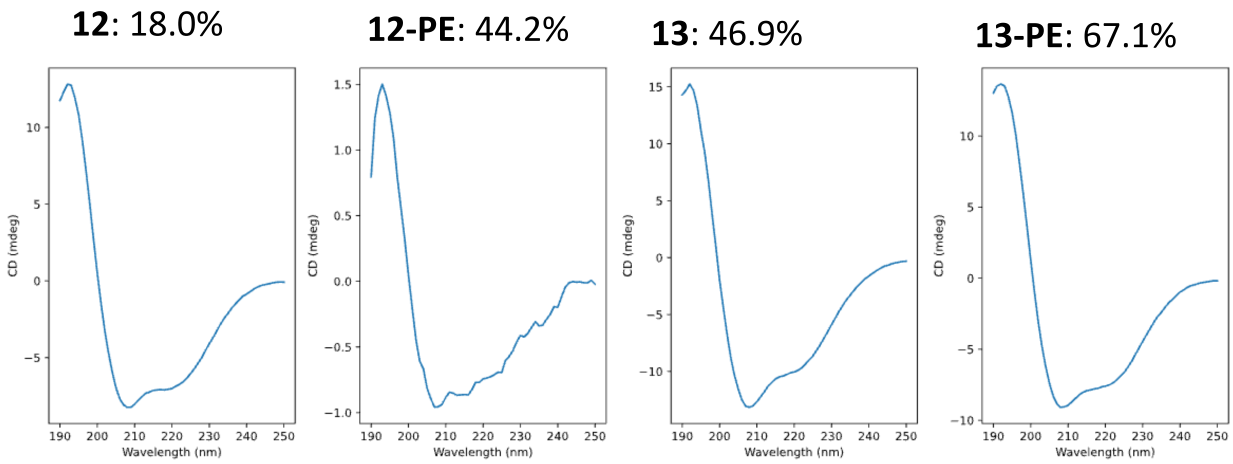


**Figure S4:** Circular dichroism measurements for the peptides chemically synthesized in this study. See Methods for information on the calculation of alpha helicity.

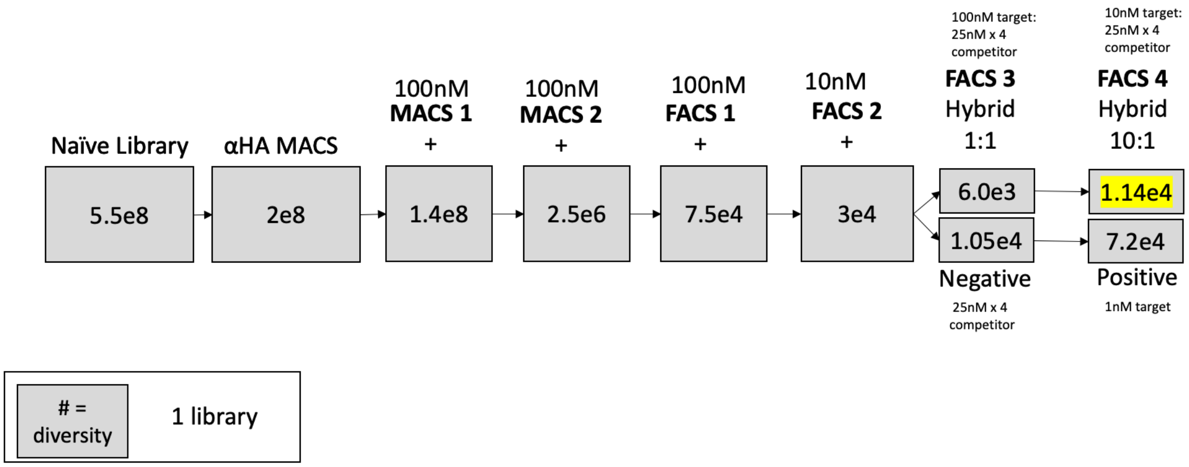


**Figure S5:** Directed evolution trajectory for Bcl-xL binding peptides. Diversity is measured as the number of transformants after each step of cell sorting (and therefore represents an upper limit for the number of unique sequences present).

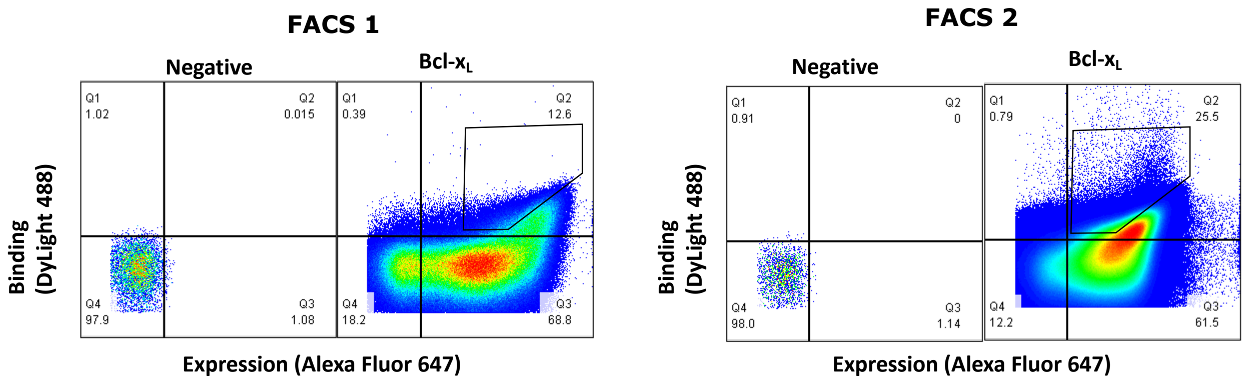


**Figure S6:** Representative flow cytometry plots for FACS 1 and 2.

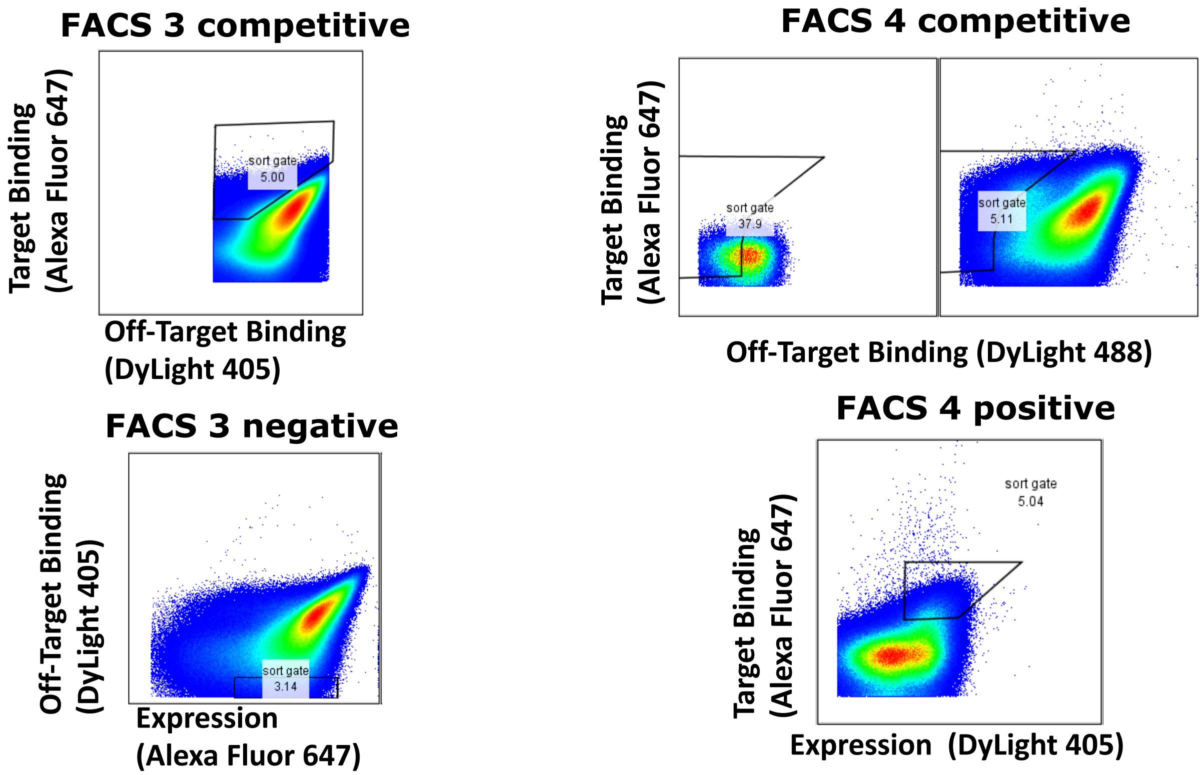


**Figure S7:** Representative flow cytometry plots for FACS 3 and 4.

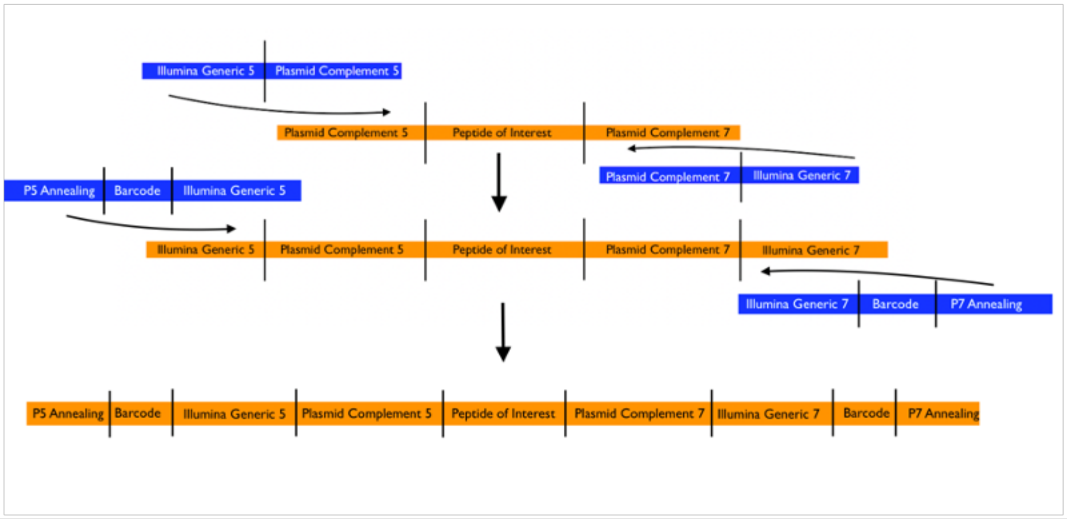


| | |
|---|---|
| 1st round forward | TCG TCG GCA GCG TCA GAT GTG TAT AAG AGA CAG CAG GTA CTT CCG TAG CTG GCC AGT CT |
| 1st reverse | GTC TCG TGG GCT CGG AGA TGT GTA TAA GAG ACA GCA CCG TAG ATG CTT GCC CAG TCG TTA |
| NGS 5-0 | AAT GAT ACG GCG ACC ACC GAG ATC TAC ACT AGA TCG CTC GTC GGC AGC GTC |
| NGS 5-1 | AAT GAT ACG GCG ACC ACC GAG ATC TAC ACC TCT CTA TTT CGT CGG CAG CGT C |
| NGS 5-2 | AAT GAT ACG GCG ACC ACC GAG ATC TAC ACT ATC CTC TGT TCG TCG GCA GCG TC |
| NGS 5-3 | AAT GAT ACG GCG ACC ACC GAG ATC TAC ACA GAG TAG ACG ATC GTC GGC AGC GTC |
| NGS 5-4 | AAT GAT ACG GCG ACC ACC GAG ATC TAC ACG TAA GGA GAT GAT CGT CGG CAG CGT C |
| NGS 5-5 | AAT GAT ACG GCG ACC ACC GAG ATC TAC ACA CTG CAT ATG CGA TCG TCG GCA GCG TC |
| NGS 5-6 | AAT GAT ACG GCG ACC ACC GAG ATC TAC ACA AGG AGT AGA GTG GTC GTC GGC AGC GTC |
| NGS 5-7 | AAT GAT ACG GCG ACC ACC GAG ATC TAC ACC TAA GCC TCC TGT GGT CGT CGG CAG CGT C |
| NGS 7-0 | CAA GCA GAA GAC GGC ATA CGA GAT TAA GGC GAG TCT CGT GGG CTC GG |
| NGS 7-1 | CAA GCA GAA GAC GGC ATA CGA GAT CGT ACT AGA GTC TCG TGG GCT CGG |
| NGS 7-2 | CAA GCA GAA GAC GGC ATA CGA GAT AGG CAG AAT CGT CTC GTG GGC TCG G |
| NGS 7-3 | CAA GCA GAA GAC GGC ATA CGA GAT TCC TGA GCC TAG TCT CGT GGG CTC GG |
| NGS 7-4 | CAA GCA GAA GAC GGC ATA CGA GAT GGA CTC CTG ATA GTC TCG TGG GCT CGG |
| NGS 7-5 | CAA GCA GAA GAC GGC ATA CGA GAT TAG GCA TGA CTC AGT CTC GTG GGC TCG G |
| NGS 7-6 | CAA GCA GAA GAC GGC ATA CGA GAT CTC TCT ACT TCT CTG TCT CGT GGG CTC GG |
| NGS 7-7 | CAA GCA GAA GAC GGC ATA CGA GAT CAG AGA GGC ACT TCT GTC TCG TGG GCT CGG |

**Figure S8:** Next generation sequencing (Illumina NovaSeq) PCR scheme and indexing primers.

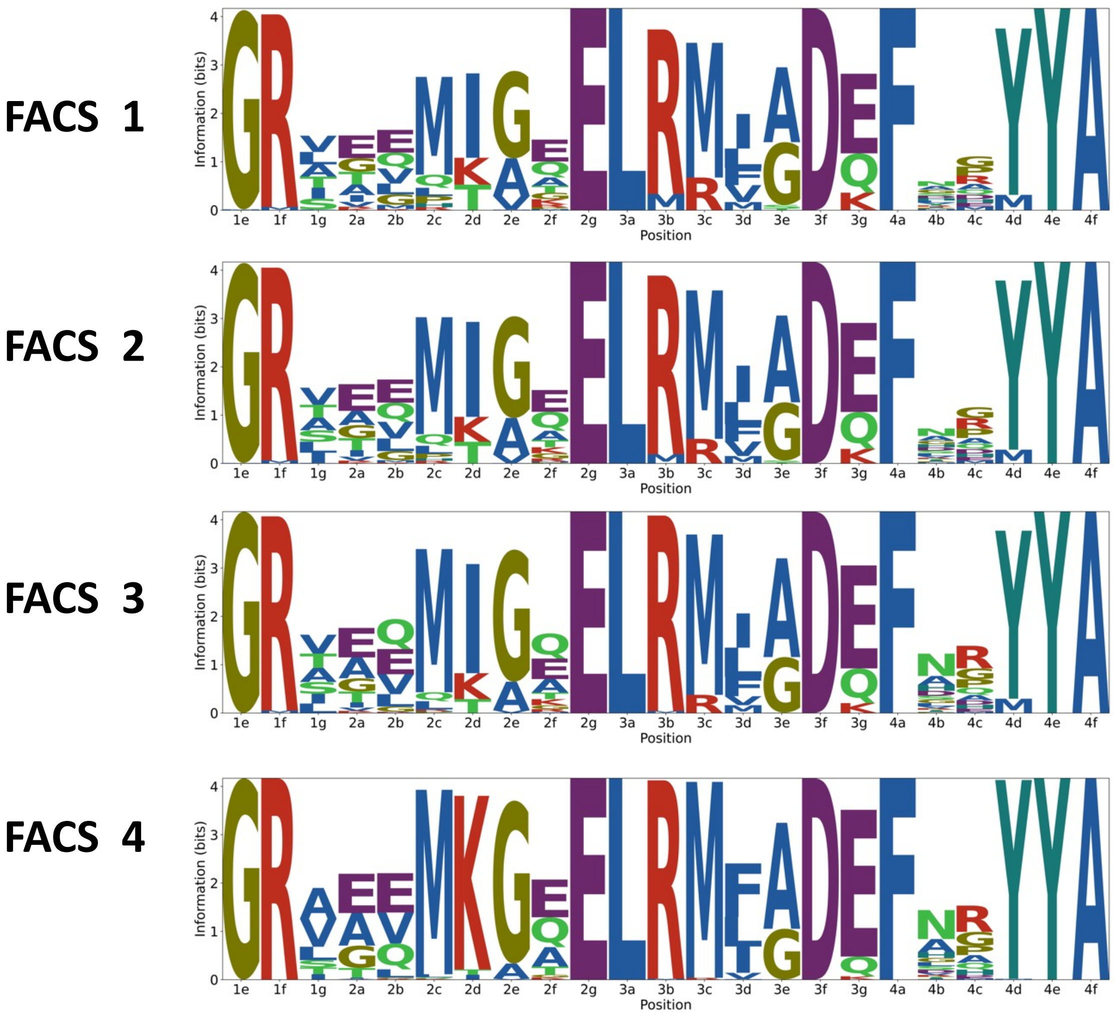


**Figure S9:** Logo plots for peptides along the directed devolution trajectory. "M" residues represent azidohomoalanine, where the peptide is stapled.

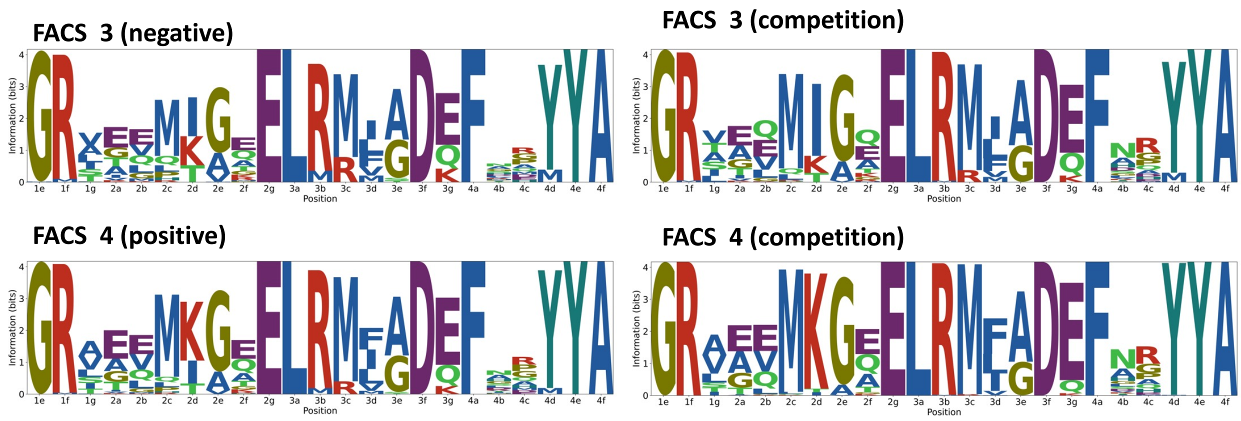


**Figure S10:** Comparison between competition methods for FACS 3 and 4 via logo plots. “M” residues represent azidohomoalanine, where the peptide is stapled.

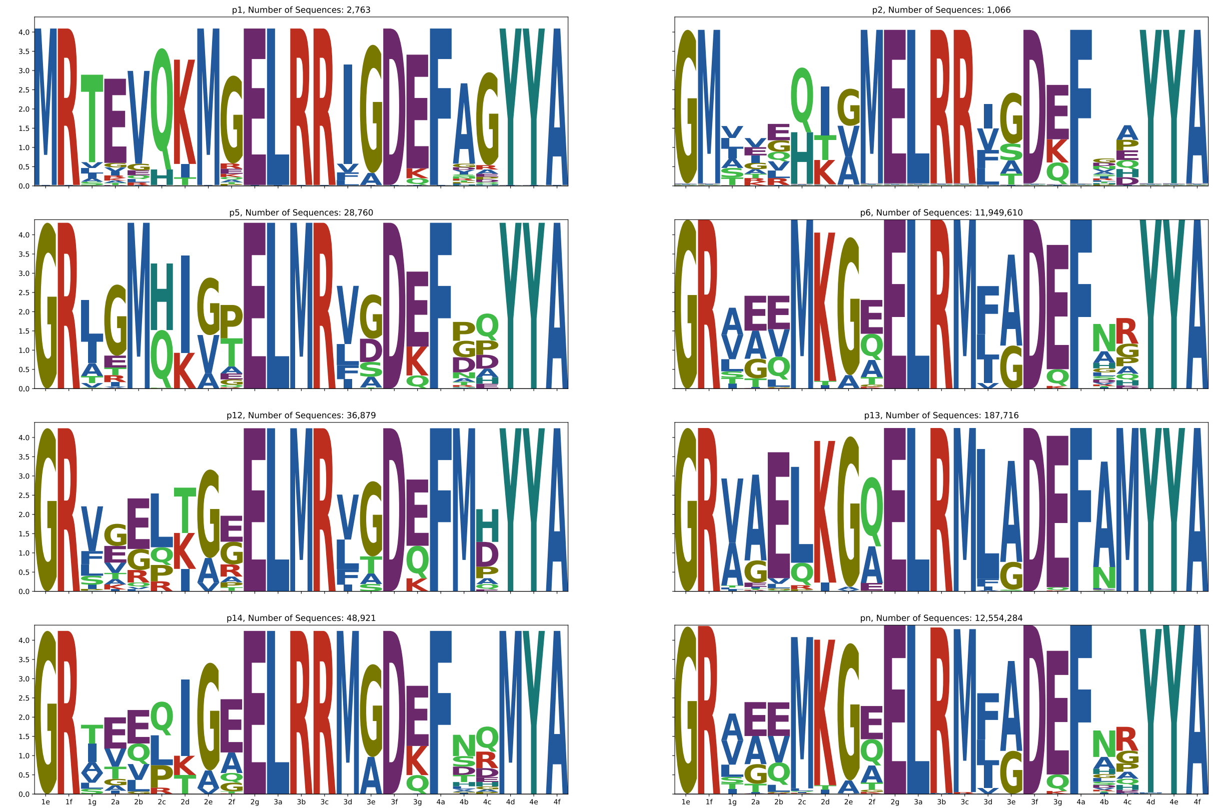


**Figure S11:** Logoplots for peptides according to their staple position (1st position: top left, 2nd position: top right, etc.).

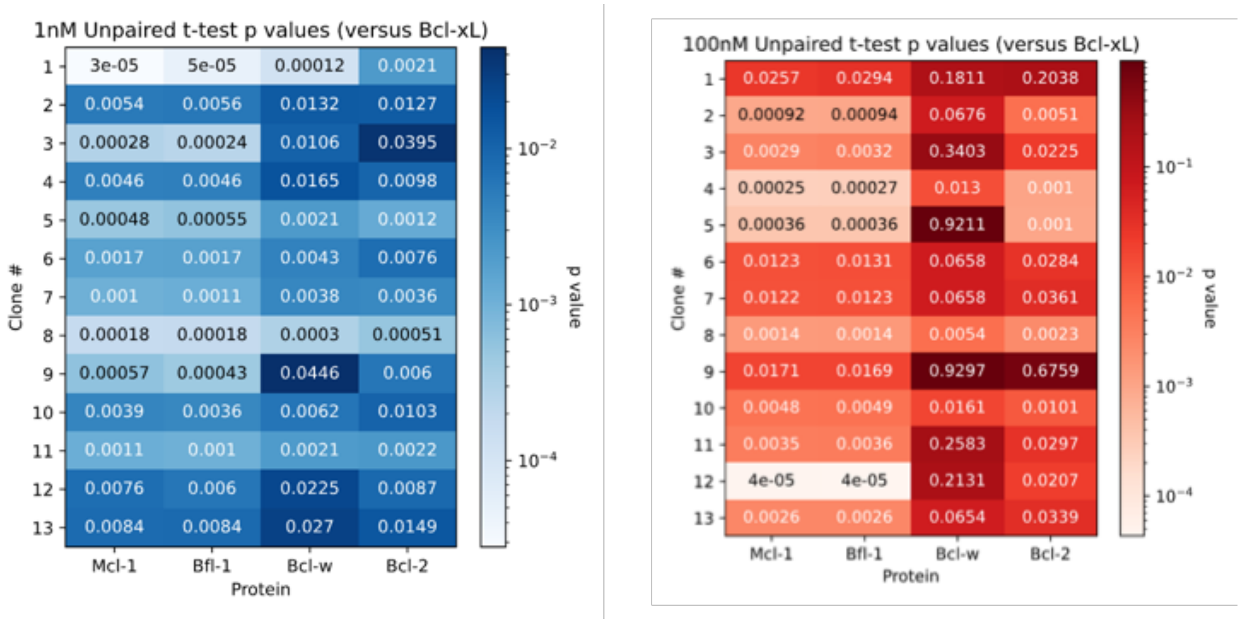


**Figure S12:** Significance of specificity for flow cytometry measurements at both [Bcl-2] = 1nM and 100nM. Significance is reported as the p-value from an unpaired t-test.

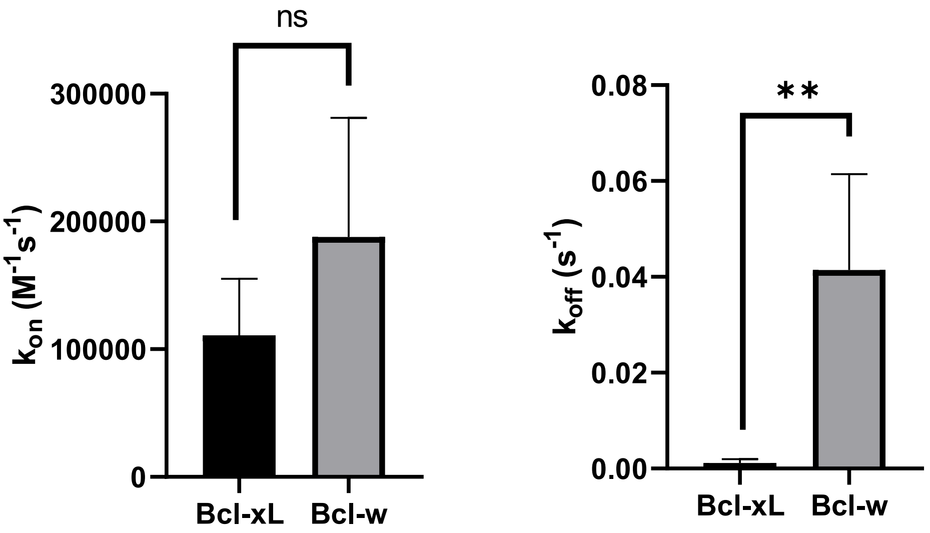


**Figure S13:** Comparison of kinetics from biolayer interferometry (BLI) data for compound **12**. The p-values for these comparisons are p=0.266 (left) and p=0.0020 (right) according to an unpaired t-test.

### Competitive Inhibition

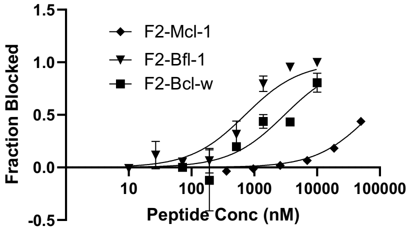


### Biolayer Interferometry

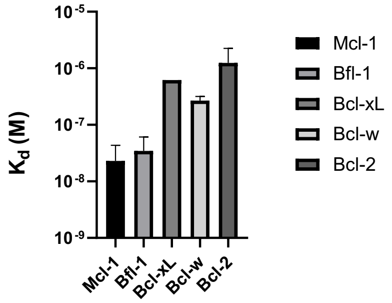


**Figure S14:** Binding affinity, measured via a competitive inhibition experiment, and mitochondrial outer membrane polarization (MOMP) for a Mcl-1 and Bfl-1 bispecific peptide (F2).
